# Systematic Engineering of Loss-of-Function Alleles in the Zebrafish Mitochondrial Proteome

**DOI:** 10.64898/2026.09.14.751483

**Authors:** Ankit Sabharwal, Md Roushan Ali, Kyler S. Mitra, Jun Morisue, Jace Klein, Rishav Sarkar, Kaila Savage, Lucy Rai Thulung, Amanda Zumbrock, Cassidy Petree, Santiago Restrepo Castillo, Mireya Mota, Gaurav K. Varshney, Karl J. Clark, Eiko Nakamuru-Ogiso, Stephen C. Ekker

## Abstract

Pathogenic variants in the 13 protein-coding genes of the mitochondrial genome underlie clinically and biochemically heterogeneous disorders. Most mtDNA-encoded genes lack defined loss-of-function (LOF) models *in vivo*. To address this gap, we have generated Z-Terminator, a systematic *in vivo* atlas of loss-of-function alleles covering all the mtDNA-encoded OXPHOS subunits in zebrafish (*Danio rerio*). We used mitochondrial TALE base editors to introduce premature termination codon (PTC) alleles via C-to-T transitions across Complexes I, III, IV, and V. Larvae harboring PTC alleles displayed bioenergetic defects and elevated lactate. While mtDNA mutations are associated with sensorineural hearing loss, the cellular basis has remained unclear. We show that engineered mtDNA LOF alleles directly impair hair cell function in proportion to heteroplasmy in a living vertebrate. We investigated the germline transmission and tissue-specific heteroplasmy of these LOF alleles and observed that a subset of variants was transmitted to the F1 generation and displayed distinct mutant loads across organs. These findings establish Z-Terminator as a vertebrate platform for interrogating the role of mtDNA protein-coding genes in cellular dysfunction and for elucidating the pathophysiology of mitochondrial disorders.

## Introduction

The human mitochondrial genome is a ∼16.6 kb compact, circular molecule governed by both nuclear and mitochondrial genetic machinery. Cells harbor multiple copies of mitochondrial DNA (mtDNA) and each molecule encodes 13 proteins that form the catalytic core of the oxidative phosphorylation (OXPHOS) complex, in addition to the 22 transfer RNAs and 2 ribosomal RNAs required for the translation of the mitochondrially encoded proteome^1–3^. Heteroplasmy, the coexistence of wild-type and mutant mtDNA copies within a cell, reflects disease penetrance and severity and contributes to phenotypic variability. Heteroplasmy can vary across tissues depending on their mitochondrial functional requirements. Individuals harboring pathogenic variants in mtDNA present with a clinically and biochemically heterogeneous spectrum of disorders, including Leber’s hereditary optic neuropathy (LHON), mitochondrial encephalopathy, and Leigh syndrome^4–7^. Mitochondrial disorders have remained largely refractory to curative treatments, primarily due to the limited ability to engineer precise edits in mtDNA, which has directly impeded the establishment of animal models.

Precise editing of mitochondrial DNA has been technically challenging, in part because inefficient exogenous RNA import into mitochondria limits the use of CRISPR-Cas systems^8,9^ and the absence of double-strand break repair^10,11^. As a result, most mechanistic studies of mtDNA disorders have relied on rho-zero^12^, cybrid cell lines^13,14^, transmitochondrial mice^15–17^, and pharmacological inhibition of mitochondrial OXPHOS complexes^18,19^. While cybrids have yielded important biochemical insights, their nuclear–cytoplasmic mismatch^13,20^ and lack of tissue-specific context limit their utility. Transmitochondrial mice carrying defined heteroplasmy have been generated, but this approach is constrained by the availability of donor cell lines and cannot be systematically extended to program pathogenic variants across all 13 mitochondrial encoded protein-coding genes. In the nuclear genome, high-throughput CRISPR screens^21–23^ and insertional mutagenesis such as gene-trap libraries^24–26^ have cataloged gene-specific contributions to the pathophysiology of nuclear-encoded mitochondrial disorders. However, no analogous resource exists for mtDNA-encoded disease, owing to the above-mentioned constraints.

Existing approaches, such as mitochondria-targeted nucleases that shift heteroplasmy by selectively degrading mutant mtDNA, cannot be used effectively to study the function of mtDNA-encoded OXPHOS subunit genes^27–29^. Mok *et al*. addressed these challenges by establishing DddA-derived cytosine base editor (DdCBE) as a CRISPR-free platform for C•G-to-T•A editing in human cells. This architecture relies on a split form of the double-stranded DNA cytosine deaminase A (DddA_tox_) fused to individual transcription activator-like effector (TALE) arms to introduce targeted C-to-T edits without generating double-strand breaks^30,31^. To facilitate the assembly of TALE base editors such as DdCBEs, our group developed the FusX TALE Base Editor (FusXTBE) platofrm, which has enabled up to 90% editing efficiency in zebrafish mtDNA *in vivo*^32^. Other groups extended the mtDNA base editing toolkit to A-to-G transitions using TALE-linked deaminases (TALEDs)^33^ and nickase-assisted tools known as mitochondrial adenine base editors (mitoABEs)^34^. More recently, unconstrained αDdCBEs that replace the conventional TALE N-terminal region with an engineered αN domain have removed the TALE 5′ - requirement, which expanded the range of targetable mtDNA sites^35^.

The mitochondrial genome represents one of the most highly conserved known contiguous stretches of vertebrate DNA between zebrafish (*Danio rerio*) and humans^36,37^. These similarities highlight the biochemical conservation of form and function between zebrafish and human mtDNA. For example, the human and zebrafish mitochondrial genomes are nearly identical in size at ∼16.6 kb and encode the same complement of 13 protein-coding genes, 22 tRNAs and 2 rRNAs. Gene order and the vertebrate mitochondrial genetic code is also fully syntenic between the two species^36,37^. Zebrafish recapitulate the tissue-specific phenotypes of respiratory chain dysfunction in a whole-organism context amenable to real-time imaging, genetic manipulation, and small molecule screening^32,36,38–43^. This vertebrate system has been employed to model with good fidelity key aspects of mitochondrial disorders caused by pathogenic variants in the nuclear-encoded mitochondrial proteins^26,44–48^.

Building on the expanding base-editing toolkit and our prior proof-of-principle demonstrating the introduction of a premature termination codon (PTC) into *mt-co3* gene locus^32^, we reasoned that the unique features of the vertebrate mitochondrial genetic code could be exploited to create systematic loss-of-function alleles. Vertebrate mitochondria use a distinct genetic code relative to the nucleus, in which the UGA codon encodes tryptophan rather than serving as a stop codon^2,37^, a feature that can be leveraged by DdCBE to introduce PTCs across mitochondrial encoded protein-coding genes. Recent work has established the feasibility of DdCBE-mediated PTC induction as an *in vivo* loss-of-function strategy for interrogating gene-specific OXPHOS dysfunction^49,50^. To date, however, a comprehensive *in vivo* loss-of-function atlas spanning all 13 mtDNA-encoded OXPHOS subunits has not been reported.

Hearing loss represents one of the most common clinical manifestations of mitochondrial dysfunction, presenting either as an isolated non-syndromic phenotype or as a feature of multisystemic disease^51–53^. Multiple pathogenic variants such as *MT-RNR1* (m.1555A>G)^54^, *MT-TL1* (m.3243A>G)^55^, and *MT-ND1* (m.3861A>C)^56^ causing hearing loss have been described across both non-coding and protein-coding mtDNA genes. Cochlear hair cells and the stria vascularis require OXPHOS to maintain the endocochlear potential and ionic gradients necessary for mechanotransduction^57^. Bioenergetic defects arising from pathogenic variants in these genes therefore are likely to affect the auditory tissue. Despite this susceptibility, the pathobiology of protein-coding mtDNA mutations in hearing loss remains understudied, largely due to the technical difficulty of generating precise, systematic loss-of-function alleles in these genes for mechanistic study.

Here we describe Z-Terminator, a systematic *in vivo* atlas of DdCBE-mediated loss-of-function alleles spanning the complete protein-coding complement of the zebrafish mitochondrial genome. We generated zebrafish mutant lines carrying PTC alleles at each locus and assessed their bioenergetic function. At representative loci, we assessed germline transmission of the mtDNA edits and measured heteroplasmy across tissues. These analyses establish heritable mtDNA loss-of-function in a vertebrate system. We further investigated the effects of mtDNA PTC alleles on lateral-line hair-cell mechanotransduction, a previously unreported consequence of mtDNA loss-of-function in a vertebrate system. All the associated metadata for these mutant lines are deposited in a publicly accessible catalog at GitHub. As a vertebrate *in vivo* resource, Z-Terminator can facilitate systematic investigations of gene-specific mitochondrial dysfunction and enable pre-clinical therapeutic studies of mtDNA-related diseases.

## Methods

### Zebrafish husbandry and ethics

Adult zebrafish and embryos were maintained according to the guidelines established by the Mayo Clinic Institutional Animal Care and Use Committee (IACUC number: A34513-13-R16), the University of Texas at Austin IACUC (IACUC number: AUP-2023-00200), the IACUC of The Children’s Hospital of Philadelphia (CHOP) (IACUC number: 23-001154), and the IACUC of the Oklahoma Medical Research Foundation (OMRF) (IACUC number: 24-28). Adult zebrafish (*Danio rerio*) of the MRF-WT background (Mayo Clinic and UT Austin), AB background (CHOP), and TAB-5 (OMRF) were maintained in a recirculating housing system at 28°C on a 14-hour light/10-hour dark cycle. Embryos were obtained by natural pairwise spawning, collected within 30 minutes of fertilization, and raised in embryo water at 28.5°C.

### Selection of premature termination codon target sites across the zebrafish mitochondrial genome

The complete coding sequences for all 13 protein-coding genes of the zebrafish mitochondrial genome were retrieved from Ensembl (GRCz11 genome build). Each gene was scanned for in-frame tryptophan (Trp) codons (TGA, read as Trp in the vertebrate mitochondrial genetic code) and in-frame glutamine (Gln) codons (CAA and CAG). In the vertebrate mitochondrial genetic code, both Trp and Gln can be converted to a stop codon TAA via a C-to-T transition on the reverse or forward strand, respectively. All 5′-TGA-3′ and 5′-TCAA-3′ motifs across the 13 genes were identified for target design. We prioritized sites where the resulting truncation removed approximately 50-60% of the protein length and where the engineered PTC was upstream of annotated functional domains. Target sites were further filtered for compatibility with TALE design rules, such as the presence or absence of a 5′ T at the N-terminal binding position for constrained versus unconstrained DdCBEs. We assessed sequence conservation at each of the predicted truncation sites between zebrafish and human orthologs by pairwise protein sequence alignment using Clustal-Omega^58^. If a prioritized site for a given protein-coding gene did not yield detectable editing (as measured via Sanger sequencing), an alternative downstream Trp or Gln codon was selected for DdCBE design, resulting in a truncation of a smaller proportion of the protein.

### Design and assembly of mitochondrial TALE base editors

Constrained^32^ and unconstrained^35^ mitochondrial TALE base editor backbone plasmids were used to generate the base editors utilized to introduce PTCs into the zebrafish mitochondrial genome. Constrained mitoTALE base editors require a 5′ T at the N-terminal TALE binding position, whereas unconstrained base editors circumvent this requirement by substituting the canonical QWS motif in the TALE N-terminal domain with an engineered RGA motif, thereby enabling targeting of sequences that lack a 5′ T^35^. Unconstrained backbones were generated from the constrained plasmid backbones pT3-FusXTBE-N and pT3-FusXTBE-C, via site-directed mutagenesis using the NEB Q5 Site-Directed Mutagenesis Kit according to the manufacturer’s instructions. In detail, the constrained backbones: pT3-FusXTBE-N and pT3-FusXTBE-C, described previously^32^ were used to assemble base editors to introduce PTCs in *mt-nd4*, *mt-nd5*, *mt-cyb*, *mt-co1*, *mt-co2*, *mt-co3*, and *mt-atp8*. Likewise, the unconstrained backbones: pT3-αFusXTBE-N and pT3-αFusXTBE-C, were used to introduce PTCs in *mt-nd1*, *mt-nd2, mt-nd3*, *mt-nd4l*, *mt-nd6*, and *mt-atp6*, where the absence of a 5′ T at the intended TALE binding site precluded the use of constrained constructs. Repeat variable di-residues (RVDs) specific to each prioritized target locus were cloned into the receiving vector backbones using the one-step FusX assembly system^32,59^. For all base editors, their corresponding spacer and RVD sequences are provided in the Supplementary Table 1. Assembled plasmids were transformed into NEB Stable Competent E. coli (High Efficiency, NEB, C3040H) cells, the cells were then subjected to blue-white screening, and positive white colonies were selected for further processing. Plasmid DNA was isolated from the positive colonies using the QIAprep Spin Miniprep Kit (QIAGEN). All constructs were verified by whole-plasmid sequencing (Plasmidsaurus, USA) prior to mRNA synthesis.

### mRNA synthesis and zebrafish microinjection

Sequence-verified plasmids were linearized using the PstI restriction enzyme and used as templates for *in vitro* transcription with the T3 mMESSAGE mMACHINE Transcription Kit (Thermo Fisher Scientific). *In vitro* transcribed mRNA was purified using the Monarch® Spin RNA Cleanup Kit (New England Biolabs), and the integrity of the purified mRNA was confirmed by agarose gel electrophoresis. Three nanoliters of mRNA at a concentration of 75–100 pg per base editor arm were injected into single-cell stage zebrafish embryos. Injected embryos were raised in embryo water at 28.5°C.

### mtDNA genotyping in zebrafish embryos

To calculate the mtDNA heteroplasmy, individual 24 hours post fertilization (hpf) embryos were transferred to PCR tubes and embryo water was aspirated. To each tube, 60 µL of 50 mM NaOH was added, and the tubes were incubated at 95°C for 30 minutes at 500 rpm in a shaking incubator, followed by the addition of 12 µL of 1 mM Tris-Cl (pH 8.0) and mixing. Lysates were used as templates for PCR amplification using NEB OneTaq 2X Master Mix as per manufacturer’s instructions, with mtDNA-specific primers listed in Supplementary Table 2. PCR amplicons were resolved on a 2% agarose gel to verify the expected amplicon size and analyzed via Sanger sequencing (GENEWIZ, Azenta Life Sciences, USA) to assess editing efficiency and detect bystander edits within the protospacer regions. Raw .ab1 files were analyzed using the EditR software^60^, which calculates mtDNA heteroplasmy from chromatogram peak heights at each target cytosine.

### Measurement of mitochondrial respiratory complex activities and lactate

Respiratory chain enzyme activities and lactate concentrations were measured in mtDNA zebrafish mutants harboring PTC alleles as described previously^32^. Twenty 7 dpf larvae per biological replicate were collected, washed two times with E3 embryo buffer, immediately frozen in liquid nitrogen, and stored at −80°C prior to downstream analysis. To prepare mitochondrial-enriched fractions, frozen larvae were homogenized on ice in a buffer consisting of 250 mM sucrose, 20 mM Tris-HCl, and 3 mM EDTA (pH 7.4) using a motorized pestle, followed by three freeze/thaw cycles. Homogenates were then subjected to differential centrifugation to obtain mitochondrial-enriched fractions. Enzyme activities were measured at 30°C in a 170 µL reaction volume using a Tecan Infinite 200 PRO microplate reader.

Complex I and Complex II activities were estimated spectrophotometrically by monitoring the reduction of 2,6-dichlorophenolindophenol (DCPIP) at 600 nm. The reaction mixture for Complex I contained 25 mM KH₂PO₄ (pH 7.4), 5 mM MgCl₂, 3 mg/mL BSA, 25 µM ubiquinone Q1, and 5 µM antimycin A. NADH (100 µM) was used to initiate the reaction, and rotenone-sensitive activity was determined by subtracting the rate measured in the presence of 5 µM rotenone. Complex II activity was assessed in the same buffer with the addition of 5 µM rotenone, using 20 mM succinate as the substrate. Complex IV activity was determined by measuring the rate of reduced cytochrome c oxidation at 550 nm in a buffer containing 25 mM KH₂PO₄ (pH 7.4), 5 mM MgCl₂, 0.015% n-dodecyl-β-D-maltoside, 5 µM antimycin A, and 5 µM rotenone. The reaction was initiated by the addition of 15 µM reduced cytochrome c, and activity was expressed as a first-order rate constant. Citrate synthase activity was measured in parallel to serve as an index of mitochondrial content. All enzyme activities were normalized to protein concentration estimated by Bradford assay^61^ or to larval number.

To measure lactate levels, frozen larvae were homogenized in 0.5 M perchloric acid on ice using a motorized pestle, followed by brief sonication and a single freeze/thaw cycle. The homogenate was centrifuged at 16,000 × g for 15 minutes, and the resulting supernatant was neutralized on ice using 1 M potassium carbonate. Following a second centrifugation at 16,000 × g for 10 minutes, the supernatant was used for downstream analysis. Lactate was measured using a colorimetric assay based on the lactate oxidase reaction, as previously described^38^. Briefly, 5 µL of extract was added to 155 µL of assay buffer containing 0.2 mM DA-64 dye (FUJIFILM Wako Chemicals), 1 mM EDTA, 0.1% Triton X-100, and 5 U/mL horseradish peroxidase in 100 mM HEPES (pH 7.4). Following a 3-minute pre-incubation at 37°C, the reaction was initiated with 10 µL of lactate oxidase (2 U/mL), and absorbance at 727 nm was recorded every 20 seconds for 15 minutes. Lactate concentrations were calculated using a sodium L-lactate standard curve and normalized per fish or per microgram of protein. All assays were carried out on six independent biological replicates, each consisting of 20 pooled 7 dpf larvae.

### Assessment of hair cell function using YO-PRO-1 vital dye uptake

To assess lateral line hair cell function, YO-PRO-1 (Quinolinium, 4-[(3-methyl-2(3H)-benzoxazolylidene)methyl]-1-[3-(trimethylammonio)propyl]-diiodide; ThermoFisher Scientific, Y3603) was used as a vital dye that selectively labels hair cells within zebrafish neuromasts^62^. The dye was reconstituted in DMSO and diluted to a working concentration of 2 µM in E3 embryo media. At 5 dpf, *mt-nd1, mt-co2, mt-cyb*, and *mt-atp8* mutants, and wild-type control larvae were washed three times in embryo media, incubated in YO-PRO-1 for 1 h at room temperature in PYREX 9-depression glass spot plates, then washed three times with embryo media. Larvae were transferred to quadrant petri dishes and maintained in embryo media under dark conditions until imaging. Imaging was performed on a Nikon Eclipse Ti2 inverted microscope equipped with a Yokogawa CSU-W1 spinning disk confocal scanning unit using 488 nm excitation. Prior to imaging, larvae were anesthetized in E3 media supplemented with 0.016% tricaine (MS-222) and immobilized in 1% low-melting-point agarose containing 0.016% tricaine to maintain anesthesia throughout image acquisition.

Fluorescence intensity was quantified using ImageJ. All images were blinded by a secondary investigator before analysis, and all measurements were performed by a single analyst. For each larva, the MI1 neuromast was identified, manually segmented, and used as a consistent anatomical reference across all specimens. The region of interest, mean gray value, and integrated density were recorded as previously described^62^. Corrected total fluorescence (CTF) was calculated by measuring background fluorescence in three regions adjacent to the MI1 neuromast, averaging these values, and applying the formula: CTF = Integrated Density − (Area of MI1 × Mean Background Fluorescence). CTF values were compared across experimental groups to assess differences in hair cell function across mitochondrial gene variants.

### Tissue-specific distribution of mtDNA heteroplasmy in adult zebrafish

To assess how mutant mtDNA was distributed across major somatic and germline tissues in adult zebrafish, mtDNA heteroplasmy was quantified in mutant females harboring edits in *mt-nd1, mt-nd5*, and *mt-atp8* genes. Mutant founder females that were confirmed positive by fin-clip genotyping were euthanized, and major organs including brain, eye, heart, liver, and ovary were dissected under a stereomicroscope. Individual tissues from each animal were processed separately to retain tissue-specific heteroplasmy profiles. Dissected tissues were stored at −80°C until further use. Total DNA was extracted from each tissue using the DNeasy Blood and Tissue Kit (QIAGEN) according to the manufacturer’s protocol. mtDNA loci encompassing the edits were PCR-amplified, and amplicons were submitted for Sanger sequencing (GENEWIZ, Azenta Life Sciences, USA). mtDNA heteroplasmy was quantified from the chromatogram files (.ab1) using the EditR software^60^.

### Acoustic-evoked behavioral responses (AEBRs) assay

Acoustic-evoked behavioral responses (AEBRs) were measured in zebrafish larvae using a DanioVision observation chamber equipped with the integrated tapping device and EthoVision XT video-tracking software (Noldus Information Technology, Wageningen, the Netherlands). At 5 dpf, individual larvae from each group (uninjected, control, and experimental) were transferred to 96-well plates containing 175 µL of E3 embryo medium. Plates were placed in an incubator and maintained at 28°C overnight. AT 6 dpf, plates were transferred to the DanioVision chamber for testing. Larvae were allowed to acclimate for 30 min before stimulus presentation. Acoustic/vibrational stimuli were delivered using the DanioVision tapping device at an intensity setting of 6. Each larva received 12 stimuli separated by a 20-s interstimulus interval. Larval movement was recorded and analyzed automatically using EthoVision XT. The percentage of responses was calculated by counting the number of responses after stimuli when there was no movement in the two seconds before the stimuli and excluding any embryos that moved before 6 or more stimuli. Results were analyzed using the Kruskal-Wallis test and plotted using Graphpad Prism.

### Germline assessment of mtDNA heteroplasmy

Stable germline transmission of mtDNA edits generated by mitochondrial TALE base editors was assessed in F0 founder females identified by fin-clip genotyping. Founder F0 females were then outcrossed to wild-type males, and F1 embryos were collected from individual crosses. mtDNA heteroplasmy was quantified in F1 progeny across edited loci in Complexes I and V. Complex I targets (*mt-nd1* and *mt-nd5*) and Complex V target (*mt-atp8*) were prioritized for germline analysis. Genomic DNA was extracted from individual F1 embryos by NaOH lysis followed by neutralization with Tris-Cl (pH 8.0). mtDNA loci spanning the edited sites were amplified by PCR, and heteroplasmy levels were determined by Sanger sequencing. Raw chromatogram files (.ab1) were analyzed using the EditR software^60^ to calculate the frequency of C-to-T transitions in the editing windows in F1 animals.

### Z-Terminator Web Resource Development

To disseminate information on the collection of zebrafish mtDNA mutants generated in this study, we built the Z-Terminator web resource, a searchable online database of characterized mutant lines. The website was designed and developed with the assistance of Google Antigravity^63^, an AI-powered agentic coding assistant. The site is deployed as a static, multi-page application built with standard web technologies (HTML, CSS, and JavaScript). The web interface includes an interactive mutant catalog that allows users to search and filter mtDNA mutants by gene name, phenotype, or editor type, and to download associated metadata including plasmid maps, annotated amplicons, and protein alignment files for each line. The full source code is publicly available on GitHub (*catalog to be released upon publication*). The website is hosted on GitHub Pages (*catalog to be released upon publication*) and automatically redeploys the resource when the repository is updated. No login or registration is required, and the resource is freely accessible to the scientific community.

### Statistical analyses

All statistical analyses were performed in GraphPad Prism 11. Differences between experimental and control groups were assessed using Welch’s unpaired t-test and Kruskal-Wallis test. Exact p-values and test details are reported in the figure legends.

## Results

### Z-Terminator targets all 13 mitochondrially encoded OXPHOS subunits in the zebrafish mitochondrial genome

Mitochondrial TALE base editors were designed targeting each of the 13 protein-coding genes of the zebrafish mitochondrial genome. Together, both constrained and unconstrained editor architectures (Figure 1A, 1B) together enabled the identification of a PTC-inducing target locus in all 13 genes (Figure 1C), covering every OXPHOS complex encoded by the mitochondrial genome. This constitutes the first comprehensive *in vivo* toolkit for loss-of-function alleles of all 13 mitochondrially-encoded OXPHOS subunits in a vertebrate model organism (Figure 2A). Initially, we designed and tested TALE 5’-T constrained base editors to introduce PTC alleles in all mitochondrial encoded protein-coding genes. However, we observed that these constrained base editors yielded detectable editing for 7 out of 13 genes (*mt-nd4, mt-nd5, mt-cyb, mt-co1, mt-co2, mt-co3, mt-atp8*). For the remaining 6 genes (*mt-nd1, mt-nd2, mt-nd3, mt-nd4l, mt-nd6, mt-atp6*), the constrained base editors failed to yield detectable editing at the prioritized target sites. The αDdCBE architecture circumvents the TALE 5’-T requirement of canonical DdCBEs^35^, allowing the TALEs to be repositioned along the target sequence for these 6 loci. This TALE shifting strategy expanded the pool of designable sites for PTC engineering at mtDNA loci that would otherwise have been inaccessible. The complete set of Z-Terminator target sites spans all of the four respiratory complexes with mitochondrially-encoded subunits, distributed across the 16,596 bp zebrafish mitochondrial genome (Figure 1C): Complex I (*mt-nd1*, m.4356G>A; *mt-nd2*, m.5350C>T; *mt-nd3*, m.10822G>A; *mt-nd4*, m.11513G>A; *mt-nd4l*, m.11148G>A; *mt-nd5*, m.13311C>T, *mt-nd6*, m.14967C>T), Complex III (*mt-cyb*, m.15729G>A), Complex IV (*mt-co1*, m.6962C>T; *mt-co2*, m.8617G>A; *mt-co3*, m.10215C>T), and Complex V (*mt-atp6*, m.9412C>T; *mt-atp8*, m.9008C>T). The RVD sequences for each target locus are provided in Supplementary Table 1.

**Figure 1:**
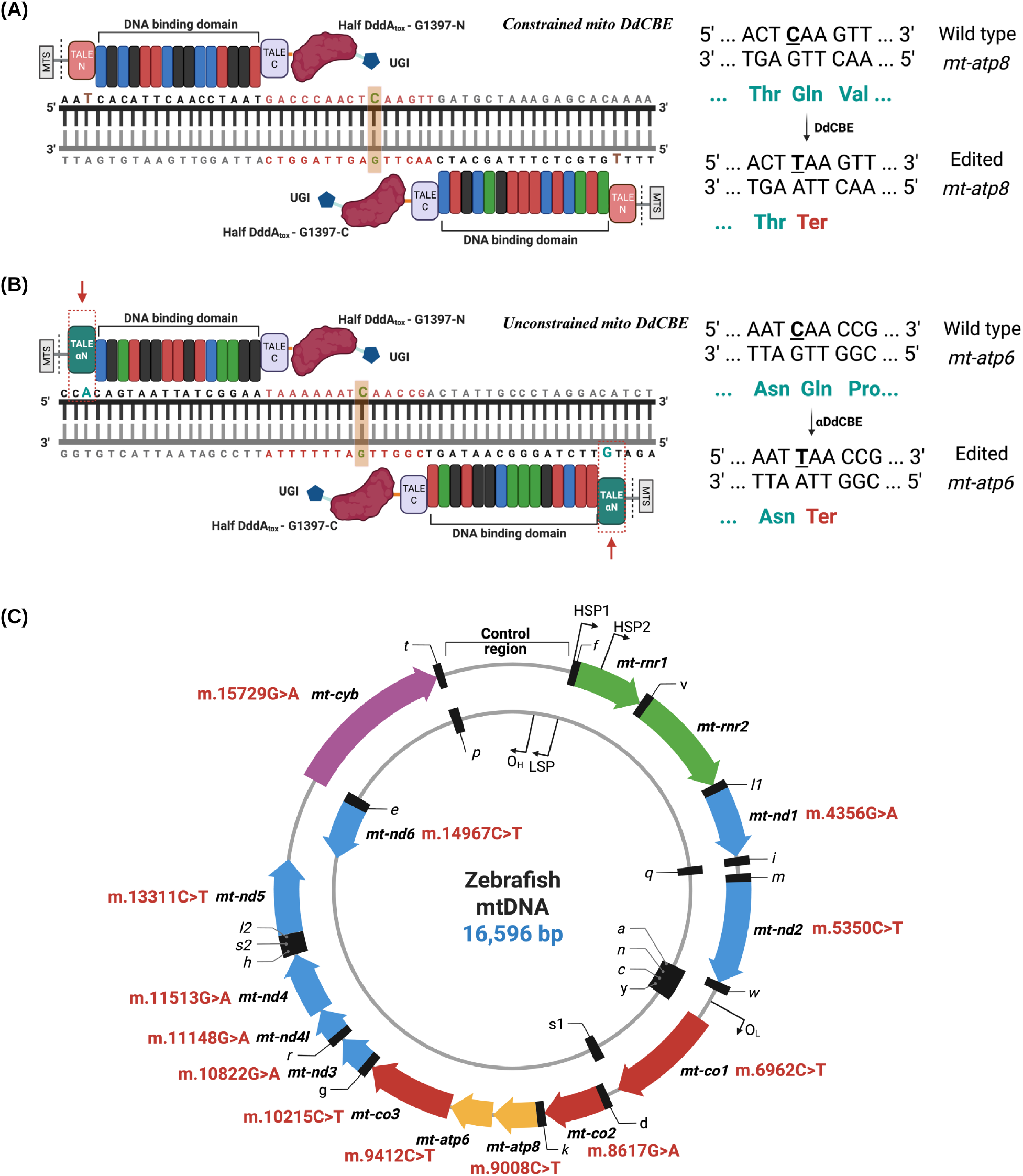
Schematic of the mitochondrial base editor architecture and engineered stop codon sites in mitochondrial DNA. **(A)** Constrained mitochondrial DdCBE architecture. Each base editor arm is fused to the Idh2 mitochondrial targeting signal (MTS) (highlighted in grey) to facilitate mitochondrial import. The repeat variable diresidues of the TALEs arms are shown in different colors (green-guanine-NN; blue-cytosine-HD; red-thymine-NG; black-adenine-NI). The spacer region, which contains the editing window, is depicted in red. As an example, a genomic target cytosine within *mt-atp8* locus is highlighted in green with bold font. The constrained DdCBE converts a CAA (Gln) codon to TAA (Ter), introducing a premature termination codon. **(B)** Unconstrained mitochondrial αDdCBE architecture. The αDdCBE replaces the standard TALE N-terminal region with an engineered TALE αN domain (shown in green and indicated by a red arrow), relaxing the 5’-T requirement of the constrained editor. αDdCBE-mediated C-to-T conversion of CAA to TAA, introducing a premature termination codon in *mt-atp6* is shown as an example. **(C)** Map of the zebrafish mitochondrial genome, with Z-Terminator target sites shown in red with their genomic coordinates. Figure 1 was created using BioRender.com.

**Figure 2:**
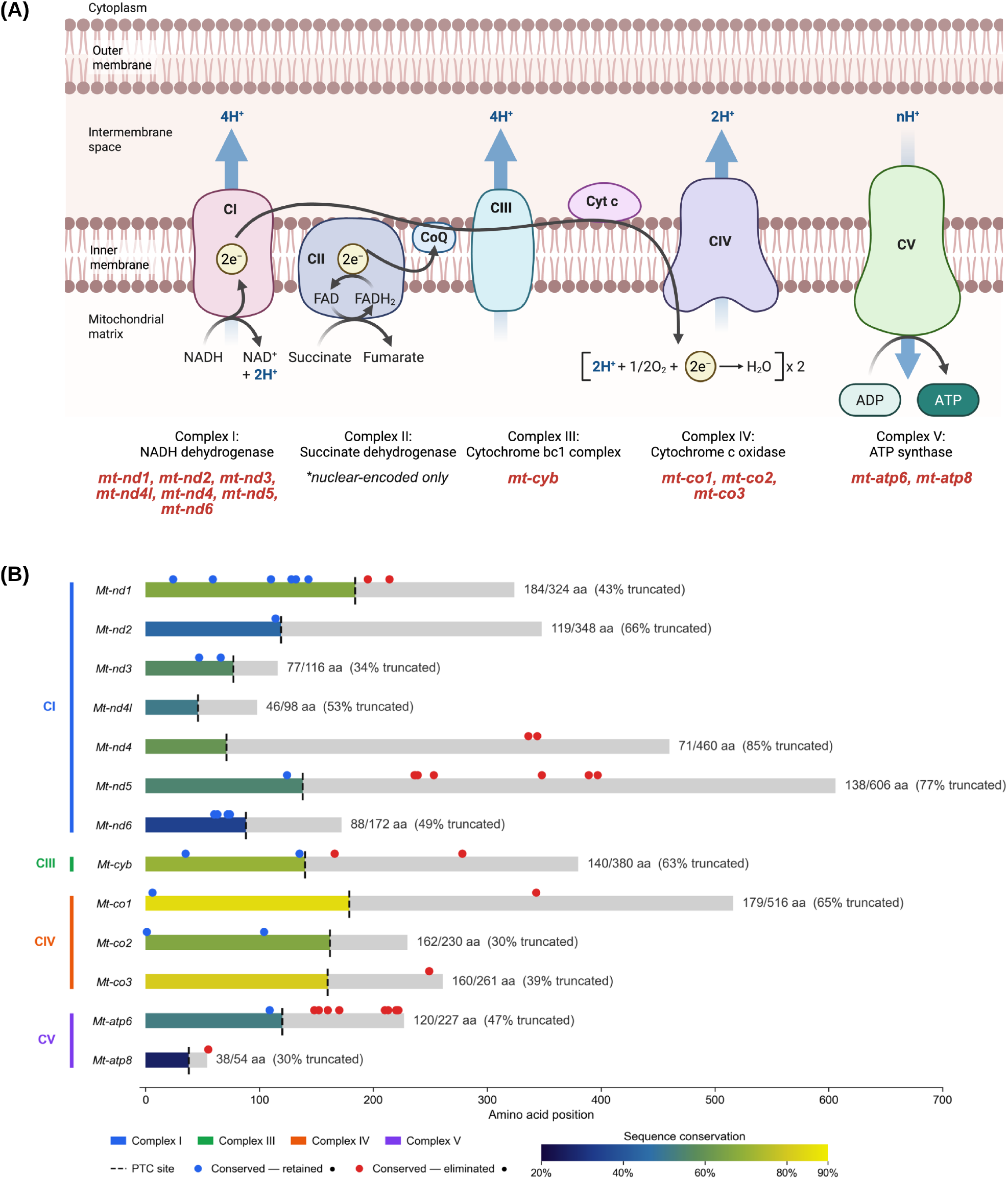
Z-Terminator introduces premature termination codons across the zebrafish mtDNA-encoded proteome. **(A)** Schematic of the mitochondrial electron transport chain representing all five OXPHOS complexes (CI-CV). mtDNA-encoded subunits are listed in red (Complex II is entirely nuclear-encoded). **(B)** Protein truncation map for all 13 zebrafish mtDNA-encoded proteins. Each bar (gray) represents the full length wild-type protein, and the color gradient shows the percentage of amino acid sequence conservation between human and zebrafish orthologs. The gradient-colored areas show the remaining protein sequence after truncation (dashed line). Conserved pathogenic variants from MITOMAP are represented by circles across the full-length protein. Variants retained in the N-terminal fragment are marked by blue circles and those within the truncated C-terminal region are shown as red circles. Retained amino acid length, and percent truncation are shown for each mtDNA-encoded protein. Figure 2A was created using BioRender.com.

To assess sequence conservation between the human gene and each zebrafish ortholog, we performed pairwise sequence alignments and mapped each target site onto the corresponding full-length zebrafish protein sequence (Figure 2B). Predicted truncations ranged from 30% of the protein for *mt-co2* and *mt-atp8* to 85% for *mt-nd4*. In addition, we examined whether pathogenic variants were conserved in the wild-type and mutant alleles. We found that across the protein-coding genes, conserved pathogenic variants were present within the truncated C-terminal regions (Figure 2B, red markers) for genes such as *mt-nd1, mt-nd4, mt-nd5, mt-cyb, mt-co1, mt-co3, mt-atp6* and *mt-atp8*. This pattern was consistent across Complex I subunits, with the largest truncations (*mt-nd4*, 85%; *mt-nd5*, 77%) as well as for Complex V subunits (*mt-atp6*, 47%). However, for genes such as *mt-nd2*, *mt-nd3*, *mt-nd6*, and *mt-co2*, the retained N-terminal fragment preserved all known conserved pathogenic variants (Figure 2B, blue markers) as compared to their corresponding truncated C-terminal domains.

### DdCBE-mediated editing generates detectable heteroplasmy at all 13 mtDNA protein-coding genes in F0 zebrafish embryos

To evaluate PTC installment via constrained and unconstrained DdCBEs *in vivo*, we injected *in vitro*-transcribed mRNA-encoded base editors into single-cell stage zebrafish embryos. To assess *in vivo* editing activity, we sequenced individual F0 embryos at 24 hpf. C•G-to-T•A transitions were detected at all 13 target loci (Figure 3A and 3B). Both the constrained and unconstrained DdCBEs resulted in measurable on-target editing in zebrafish mtDNA. Consistent with prior published observations^30,31^, DdCBE activity exhibited locus-dependent variability, which may reflect DNA sequence-dependent accessibility constraints for these protein-based editors (Figure 3A and 3B). Of note, constrained editor designs were initially tested for all 13 loci, successfully installing PTCs for *mt-nd4, mt-nd5, mt-cyb, mt-co1, mt-co2, mt-co3*, and *mt-atp8* only. For *mt-nd1, mt-nd2, mt-nd3, mt-nd4l*, *mt-nd6*, and *mt-atp6*, the unconstrained base editor architecture was adopted. The choice of editor architecture did not affect the editing outcome, as both designs achieved comparable heteroplasmy levels. Median heteroplasmy across the targeted loci ranged from approximately 20% at the lower range to more than 70% at the upper range (Figure 3A). Editors for *mt-nd1*, *mt-nd3*, *mt-cyb*, *mt-co1* and *mt-co3* showed the highest editing efficiency, whereas those for *mt-nd2*, and *mt-nd4l* were comparatively lower. Sanger sequencing chromatograms from control and injected embryos confirmed on-target editing in the editing windows for the target mtDNA protein-coding genes (Figure 3B). In *mt-nd4*, a bystander C-to-T transition was observed two codons downstream of the PTC site (**C**TT→**T**TT; Leu→Phe) within the editing window (Figure 3B). This bystander edit confirms to the 5′TC sequence preference of the cytosine deaminase DddA, and similar bystander edits within the editing window have been reported for DdCBEs^30–32^.

**Figure 3:**
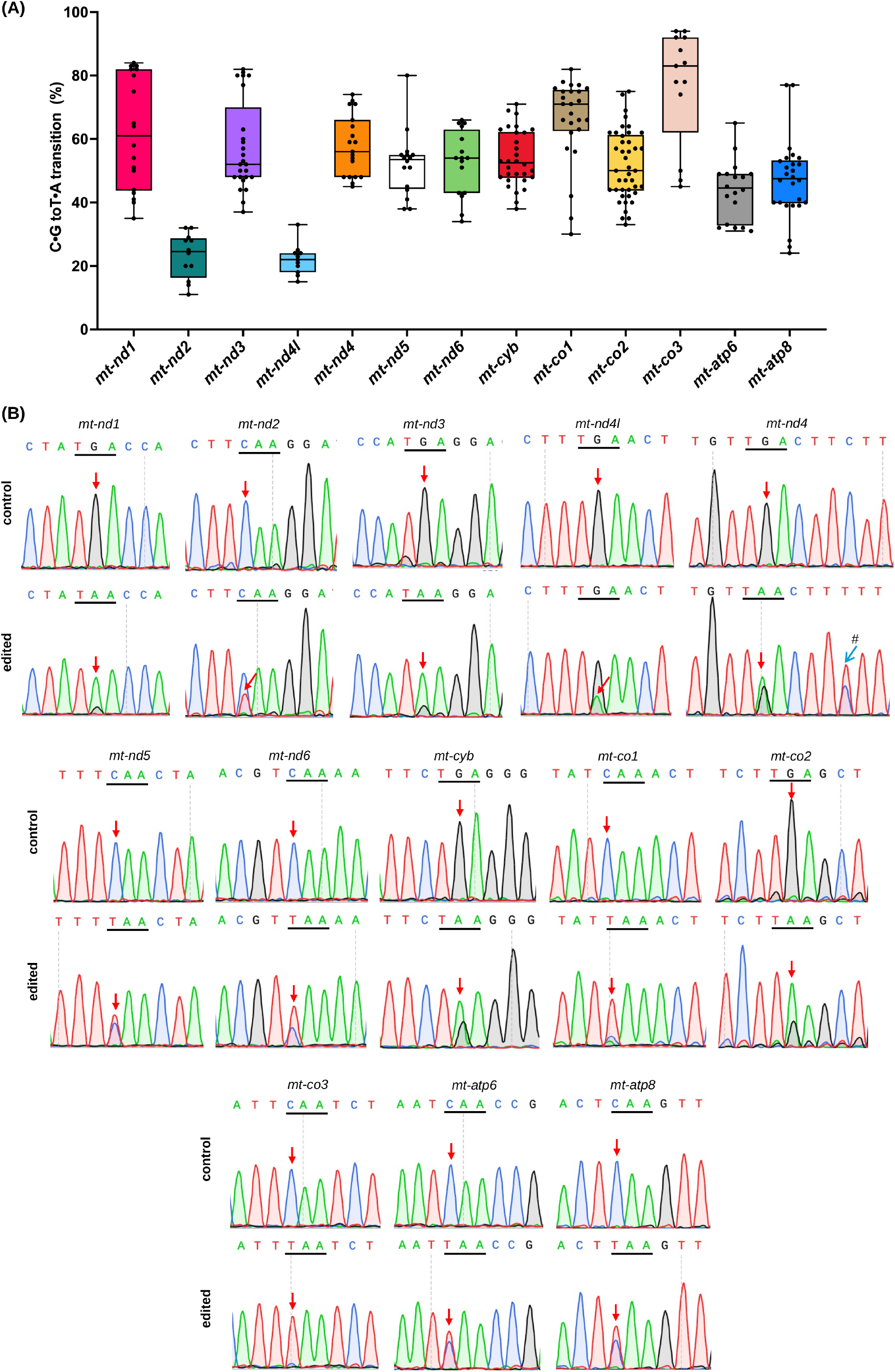
DdCBE-mediated base editing efficiency in F0 zebrafish embryos. **(A)** Box-and-whisker plots of C•G-to-T•A editing efficiencies for the favorable 5′-TC sites present in the spacer region. Data are from injected zebrafish embryos for each of the 13 mtDNA-encoded protein-coding genes. Each data point represents an individual zebrafish embryo. Editing efficiencies were calculated from the Sanger chromatograms using EditR. **(B)** Representative Sanger chromatograms of the mutant and wild-type control embryos. The target codon is underlined, and the red arrows highlight the edited cytosine residue. In *mt-nd4*, a bystander edit (#) converts CTT (Leu) to TTT (Phe) downstream of the PTC. For *mt-nd6*, chromatograms are from the reverse strand, where TCA corresponds to a TGA (termination codon) on the coding strand.

### Engineered premature termination codons impair mitochondrial bioenergetics in zebrafish larvae

To assess the functional impact of Z-Terminator alleles on mitochondrial bioenergetics, we measured respiratory chain complex activities and lactate levels in 7 dpf zebrafish larvae harboring PTCs across select mtDNA loci, encoding subunits of Complexes I, III, IV, and V. Relative to wild-type controls, Complex I activity was significantly reduced in larvae harboring PTCs in *mt-nd1, mt-nd3, mt-nd6, mt-cyb, mt-co1, mt-co2, mt-atp6*, and *mt-atp8* compared with wild-type controls (Figure 4A). This reduction varied across loci, with *mt-atp6* and *mt-co2* mutants showing the most severe deficits, and *mt-nd1*, *mt-cyb*, and *mt-atp8* mutants showing only a mild decrease. Larvae with PTCs in subunits of Complexes III, IV, and V also displayed reduced Complex I activity. This is consistent with mitochondrial supercomplex assembly, in which individual OXPHOS complexes depend on one another for stability and function^64–66^. Complex II activity was similar across almost all conditions in larvae harboring PTCs in the evaluated loci, except for a slight decrease in *mt-co1* mutants (Figure 4B). As all four subunits of Complex II are encoded exclusively by the nuclear genome, Complex II activity serves as an internal specificity control for mtDNA-directed editing, as we do not expect it to be perturbed by the Z-Terminator system. In addition, Complex IV activity was significantly decreased in larvae harboring PTCs in *mt-co1* and *mt-co2* genetic loci, both of which encode core catalytic subunits of cytochrome c oxidase (Figure 4C). Larvae with PTCs in *mt-nd3, mt-atp6* and *mt-atp8* genes also displayed reduction in the enzymatic activity of Complex IV, which can be explained by their role in supercomplex assembly. Notably, *mt-nd6* mutants displayed a slight but significant increase in Complex IV activity, potentially reflecting compensatory remodeling of the electron transport chain in response to upstream Complex I impairment. Furthermore, no significant difference in citrate synthase activity was observed across the measured genotypes (Figure 4D), except for *mt-atp8* mutants which showed a significant reduction in citrate synthase activity. In patients with mitochondrial disorders, lactate is one of the elevated clinical metabolites^67^. Impairment of mitochondrial bioenergetic function shifts cellular metabolism toward anaerobic glycolysis, resulting in the accumulation of lactate and pyruvate. Concordant with reduced OXPHOS, lactate levels were observed to be significantly altered in a subset of zebrafish mutants. Relative to wild-type controls, lactate levels were significantly elevated in *mt-nd3*, *mt-nd6*, *mt-co1*, *mt-co2*, *mt-atp6*, and *mt-atp8* mutant larvae as compared to wild type controls (Figure 4E), suggesting a metabolic shift toward anaerobic glycolysis in response to impaired OXPHOS.

**Figure 4:**
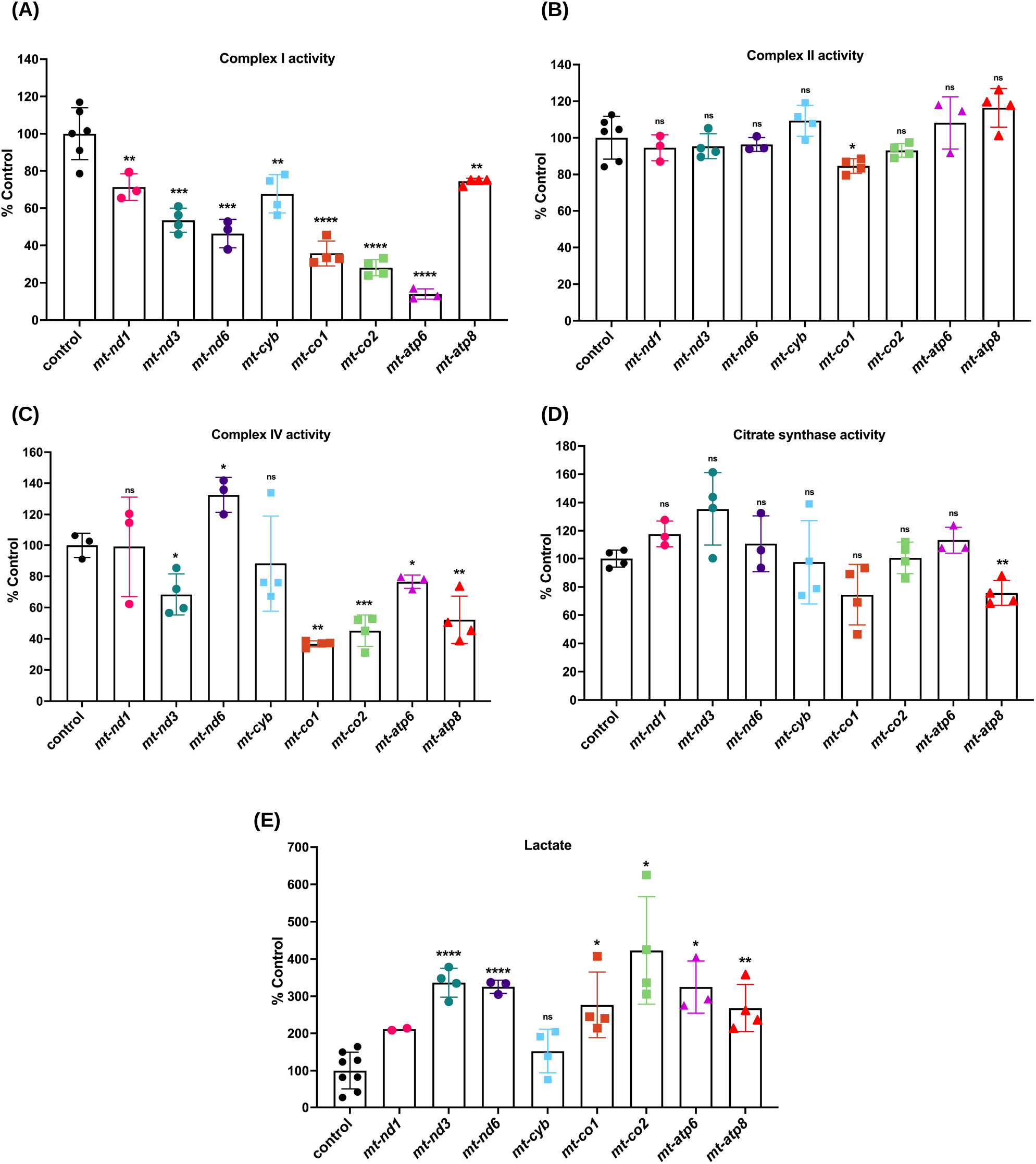
Respiratory chain complex activities and lactate levels in zebrafish mtDNA mutants. **(A-D)** Mitochondrial respiratory chain complex activities and citrate synthase activities were measured in 7 dpf zebrafish larvae. **(A)** Complex I activity was significantly reduced in larvae harboring PTC edits in *mt-nd3, mt-nd6, mt-co1, mt-co2*, and *mt-atp6*, whereas *mt-nd1*, *mt-cyb* and *mt-atp8* mutants displayed a moderate but significant decrease. **(B)** Complex II activity was maintained across all groups, except for a slight decrease in the *mt-co1* mutants. **(C)** Complex IV activity was significantly reduced *in mt-nd3, mt-co1, mt-co2, mt-atp6*, and *mt-atp8* and was slightly increased in the *mt-nd6* mutants compared with controls. No significant change was observed in the *mt-nd1* and *mt-cyb* larvae. **(D)** No difference in the citrate synthase activity was observed between the injected and control larvae across all groups except *mt-atp8*. **(E)** Lactate levels were significantly elevated in the *mt-nd3, mt-nd6, mt-co1, mt-co2, mt-atp6*, and *mt-atp8* mutants, consistent with impaired OXPHOS. Each data point represents one biological replicate consisting of 20 larvae. (*p < 0.05; **p < 0.01; ***p < 0.001; ****p<0.0001). *p*-values were determined by Student’s t-test. Error bars represent the standard deviation. dpf: days postfertilization.

### Heteroplasmy-dependent impairment of lateral line hair cell function in DdCBE-edited zebrafish larvae

Having established that Z-Terminator alleles compromise mitochondrial bioenergetics, we next examined whether these deficits extend to a functional sensory phenotype. Sensorineural hearing loss is among the most common clinical manifestations of mitochondrial disease^51,68^. Lateral line hair cells are metabolically active sensory receptors and are sensitive to mitochondrial dysfunction^69^. In zebrafish, lateral line hair cells are functionally analogous to hair cells of the mammalian inner ear^69,70^. We assessed mechanotransduction in lateral line hair cells using the YO-PRO-1 vital dye uptake assay, in which the cationic dye enters hair cells through open mechanotransduction channels^62,71^. We used the MI1 neuromast as described previously^62^ as a consistent anatomical reference across all specimens. The corrected total fluorescence (CTF) at the MI1 neuromast was calculated as an indirect readout of hair-cell functional integrity. We quantitatively assessed dye uptake in germline mutant larvae harboring PTCs in *mt-cyb*, *mt-co2*, and *mt-atp8*, each compared with wild-type controls (Figure 5A-B, 5C-D, and 5E-F). The CTF was significantly reduced in all three mutant lines relative to wild-type controls (p < 0.0001; Figure 5B, 5D, and 5F), and representative images showed a marked decrease in neuromast labeling in mutant larvae (Figure 5A, 5C, and 5E). Impaired dye uptake was thus observed across PTC alleles affecting three distinct respiratory complexes, Complex III, Complex IV, and Complex V. To determine whether the degree of hair cell impairment scaled with mutational burden, we correlated per-larva CTF with mtDNA heteroplasmy. Dye uptake was inversely correlated with heteroplasmy in all three lines, with progressively lower fluorescence at higher fractions of mutant mtDNA. For *mtDNA* mutants, heteroplasmy-dependent decline was observed suggesting that the severity of hair-cell dysfunction is proportional to the fraction of mutant mtDNA alleles.

**Figure 5:**
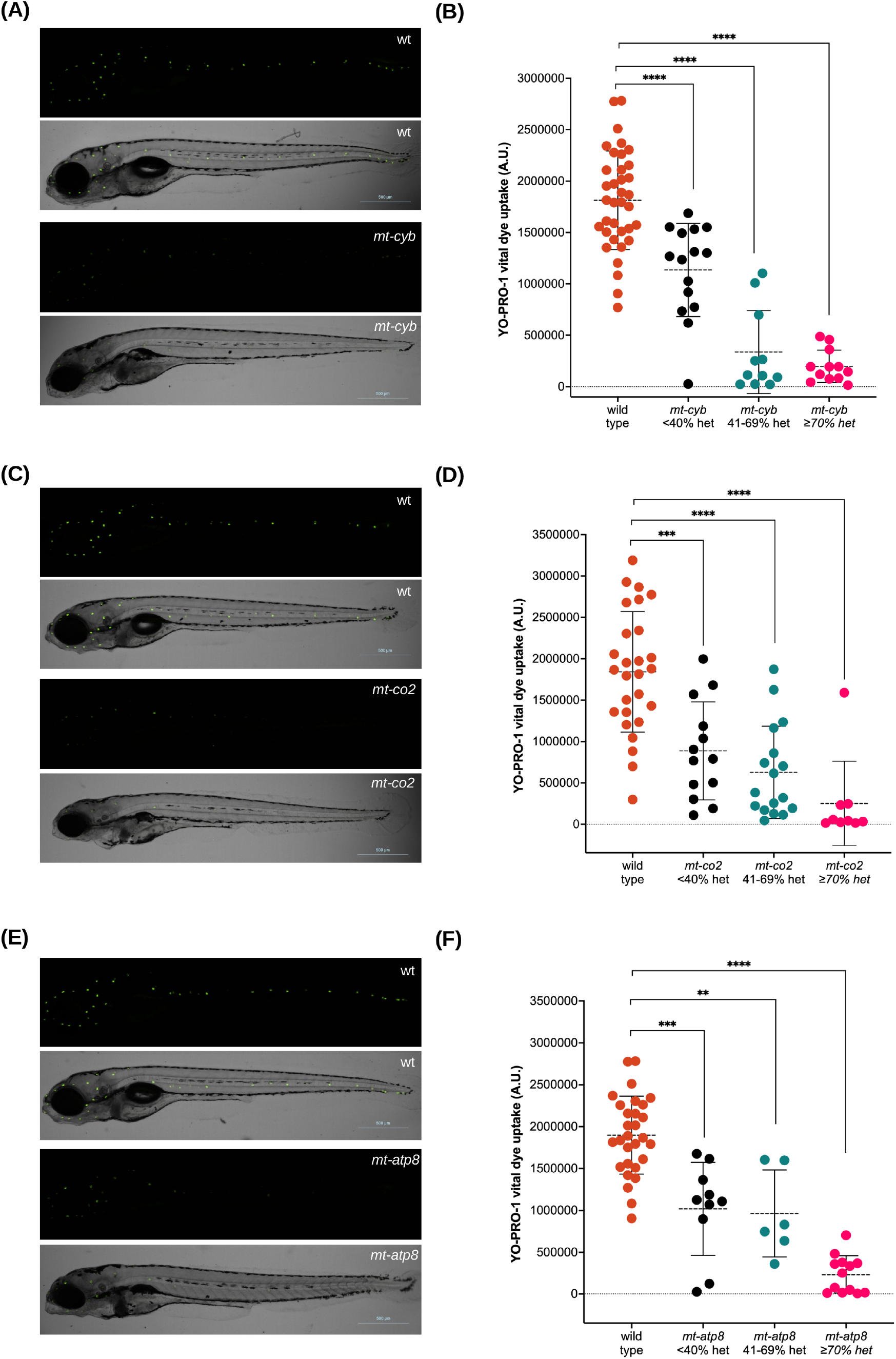
Mitochondrial premature termination codon alleles impair lateral-line hair cell mechanotransduction in a heteroplasmy-dependent manner. **(A, C, E)** Representative YO-PRO-1 fluorescence and corresponding brightfield images of wild-type and mutant larvae at 5 dpf for *mt-cyb* (A), *mt-co2* (C), and *mt-atp8* (E). Mutant larvae show a marked reduction in neuromast YO-PRO-1 labeling. Magnification-4X; Scale-bar: 500 µm **(B, D, F)** Corrected total fluorescence (CTF) of YO-PRO-1 vital dye uptake at the MI1 neuromast in wild-type versus mutant larvae displaying different mtDNA heteroplasmy for *mt-cyb* (B), *mt-co2* (D), and *mt-atp8* (F). Each point is an individual larva, and mutant points are colored by mtDNA heteroplasmy bin (see legend). p-values for wild-type versus mutant comparisons were determined by Student’s t-test (*p < 0.05; **p < 0.01; ***p < 0.001; ****p < 0.0001). Horizontal lines indicate the mean and error bars represent the standard deviation. N: *mt-cyb*, 36 wild-type and 38 mutant larvae across five biological replicates; *mt-co2*, 27 wild-type and 39 mutant larvae across four biological replicates; *mt-atp8*, 29 wild-type and 29 mutant larvae across four biological replicates.

To exclude the possibility that diminished dye uptake arose from the microinjection procedure rather than the engineered mtDNA edit, we performed an independent set of experiments in F0-injected larvae, comparing wild-type, injected-control, and mutant groups for *mt-nd1*, *mt-cyb*, and *mt-co2* (Supplementary Figure 1). Injected controls received mRNA encoding the active N-terminal base editor left arm paired with a scrambled TALE right arm. Across all three genes, *mt-nd1*, *mt-cyb*, *mt-co2* mutant larvae exhibited significantly lower CTF than injected controls (Supplementary Figure 1). Injected *mt-nd1* mutant larvae showed reduced dye uptake relative to wild-type controls. For *mt-cyb*, injected controls showed a smaller reduction relative to wild-type larvae. Nonetheless, mutant larvae remained significantly below injected controls in both cases. These results indicate that the loss of hair-cell function is specifically attributable to mtDNA editing and not to the microinjection procedure.

To determine whether mitochondrial base editing affects auditory function, we measured the acoustic evoked behavioral response (AEBR) in larvae injected with the *mt-cyb* base editor, alongside control-injected and uninjected tab5 (wild type) animals (Supplementary Figure 2). Injection per se had no detectable effect on hearing as control-injected larvae responded at 79.5 ± 4.7% (mean ± SEM, n = 26), indistinguishable from uninjected tab5 larvae (84.6 ± 4.3%, n = 26; Dunn’s post hoc after Kruskal-Wallis, p > 0.99). By contrast, larvae injected with the *mt-cyb* base editor showed a marked reduction in AEBR response rate (55.1 ± 7.6%, n = 18), significantly lower than both control-injected and uninjected tab5 larvae (p = 0.003; Kruskal-Wallis H = 11.90, p = 0.0026) (Supplementary Figure 2).

### Tissue-specific heteroplasmy distribution in DdCBE-edited F0 adult zebrafish

We next examined how DdCBE edits were distributed across metabolically diverse somatic and germline tissues in adult F0 females harboring mutations in *mt-nd1, mt-nd5*, and *mt-atp8* (Figure 6A, 6B and 6C). mtDNA heteroplasmy was broadly detected across all six analyzed tissues (brain, eye, heart, liver, caudal fin, and ovaries). This confirmed that edited alleles were systemically retained in adult F0 fish. The caudal fin-clip method was used to identify the founder females. Heteroplasmy varied across tissues within individual fish at all three loci. Brain, eye, and heart showed similar heteroplasmy within individual fish across all three genes. In *mt-nd1* mutants, tissue heteroplasmy ranged from approximately 3% to 68% across individuals (Figure 6A). Within a given individual, the somatic tissues consistently exhibited higher heteroplasmy levels compared to the ovaries. However, a subset of females maintained uniform heteroplasmy levels across all sampled tissues. Majority of *mt-nd5* F0 females displayed heteroplasmy across all six tissues. Each fish exhibited a distinct inter-individual heteroplasmy profile, spanning approximately 2% to 56% (Figure 6B). In most animals, somatic tissues showed consistent mutant mtDNA levels, whereas ovarian heteroplasmy diverged from the somatic average. The *mt-atp8* locus showed the greatest variability across the analyzed tissues, with some founder females exhibiting larger tissue-to-tissue shift than that observed for either of the Complex I genes (Figure 6C). Across all three genes, for most founder females, ovarian mtDNA heteroplasmy was lower than somatic heteroplasmy within the same individual.

**Figure 6:**
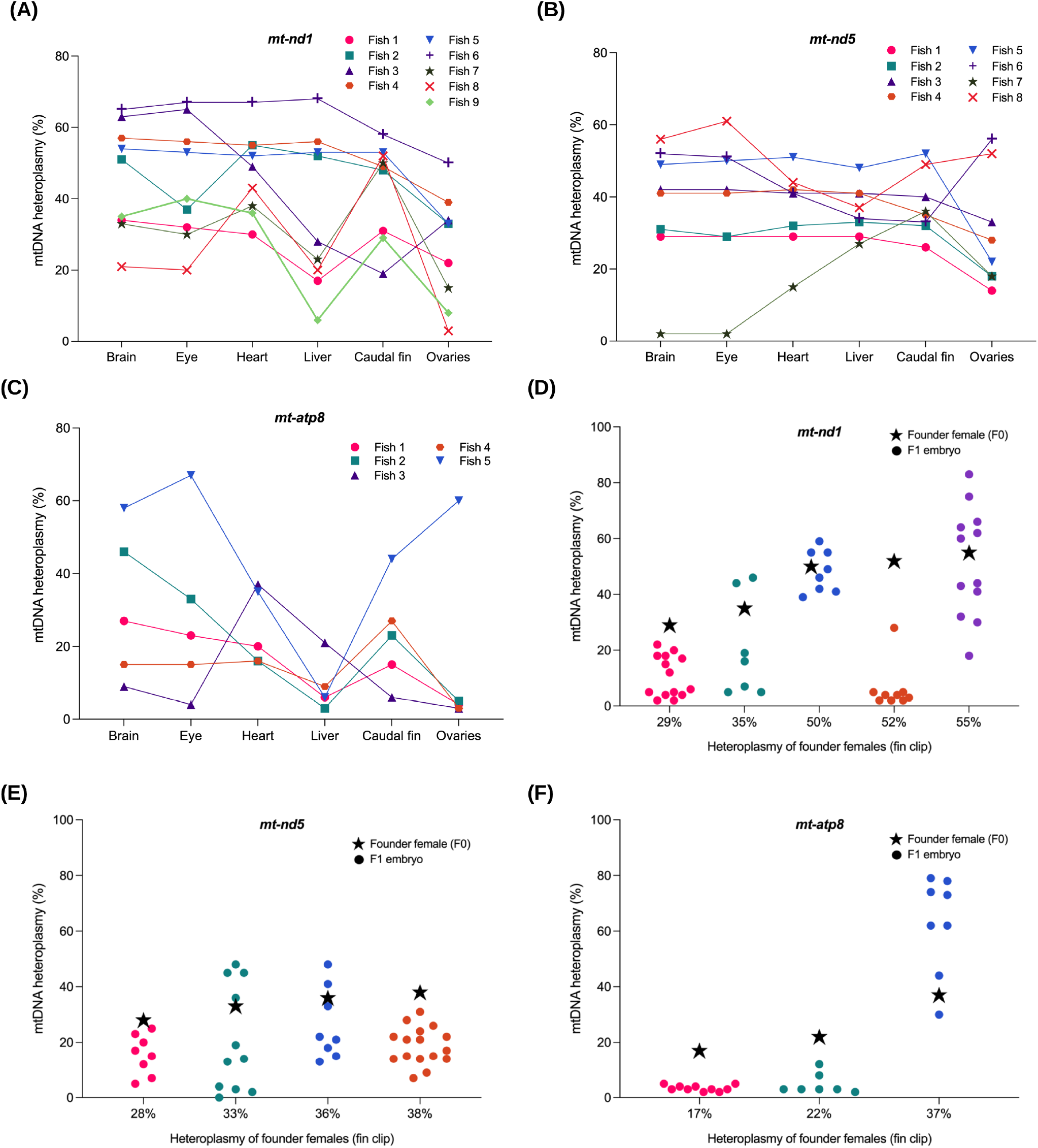
Tissue-specific heteroplasmy maintenance and germline transmission of mtDNA edits in zebrafish. **(A-C)** Tissue-specific mtDNA heteroplasmy for *mt-nd1* **(A)***, mt-nd5* **(B)**, and *mt-atp8* **(C)** female mutants. Each line connects the variation in the mtDNA heteroplasmy for the individual F0 founder females across six tissues: brain, eye, heart, liver, caudal fin, and ovaries. Each fish is distinguished by color and symbol. **(D-F)** Germline transmission of mtDNA of *mt-nd1* **(D)***, mt-nd5* **(E)**, and *mt-atp8* **(F)** edits from F0 founder females to F1 embryos. Diamond symbol denotes the heteroplasmy of the F0 founder female as measured from fin-clip. The x-axis shows the percentage of heteroplasmy for the founder females. Circles represent the heteroplasmy of the individual embryos from the outcross with wild type males. Each founder female and corresponding clutch is denoted by different color.

### Germline transmission of DdCBE-induced mtDNA mutations and heteroplasmy segregation in F1 progeny

To investigate whether edited mtDNA alleles could be propagated across generations, fin-clip–positive F0 females were outcrossed to wild-type males, and heteroplasmy was measured in individual F1 embryos at the targeted loci within *mt-nd1*, *mt-nd5*, and *mt-atp8* loci (Figure 6D, 6E, and 6F). Mutant mtDNA was detected in F1 progeny from multiple founder females across all three loci. For *mt-nd1* founder females, fin-clip heteroplasmy ranged from 29% to 55% (Figure 6D). The founder with 55% heteroplasmy transmitted edits to F1 embryos, with their heteroplasmy ranging from approximately 18% to 83%. The founders with 29% and 35% heteroplasmy produced clutches in which most embryos carried less than 20% mutant mtDNA. Two founder females with comparable somatic heteroplasmy of 50% and 52% showed contrasting patterns of germline transmission. Former transmitted edits to all sampled embryos at comparable levels of approximately 50%, while the later yielded progeny with heteroplasmy below 5%, except for a single outlier. For *mt-nd5* founder females (28%, 33%, 36%, and 38% somatic heteroplasmy), F1 heteroplasmy levels were distributed, and in several embryos deviated substantially from the maternal fin-clip value (Figure 6E). Variation in germline inheritance was most pronounced in *mt-atp8* founder females (Figure 6F). The founder with 37% somatic heteroplasmy produced a clutch in which F1 embryos reached mutant mtDNA levels as high as 80%. This exceeded the maternal somatic heteroplasmy by approximately twofold. In contrast, founders with 17% and 22% somatic heteroplasmy produced clutches in which most F1 embryos showed less than 10% heteroplasmy. The wide inter-embryo variance within the progeny of a single founder reflects a stochastic mtDNA bottleneck during oogenesis^72–74^. A restricted subset of mtDNA molecules is transmitted to each oocyte, causing rapid heteroplasmy segregation across offspring. Therefore, individual F1 siblings from the same founder can display a wide heteroplasmy range, from near-zero to well above the maternal value.

### The Z-Terminator resource: Publicly accessible mitochondrial mutant repository

No comprehensive collection of loss-of-function animal models exists for the 13 mtDNA-encoded subunits of the OXPHOS machinery. We established the Z-Terminator library to accelerate research in mitochondrial biology and make these models and their associated editing reagents freely available to the research community. The Z-Terminator resource is hosted as a web-based database that contains metadata for all 13 mutant lines generated in this study (Figure 7). The database includes genotypic and molecular metadata for each mutant line, as well as functional annotations. Metadata entries include genomic coordinates, protein alignments of zebrafish mtDNA protein-coding genes with their human orthologs, the engineered PTC site for each gene, reference and alternate alleles, the resulting amino acid change and predicted protein truncation percentage, heteroplasmy levels in F0 injected embryos, plasmid maps for the TALE cytosine base editor constructs, primer sequences for genotyping, and amplicon sizes. Sanger chromatograms (.ab1 files) are included for independent verification of editing outcomes. Users can search the database by gene name or nucleotide position and download all associated metadata for each mutant line. The academic community can request mutant zebrafish lines, husbandry protocols, and genotyping procedures through a dedicated “Contact us” tab. Open access to Z-Terminator enables direct translation from targeted mtDNA genotypes to functional analysis in a vertebrate system.

**Figure 7:**
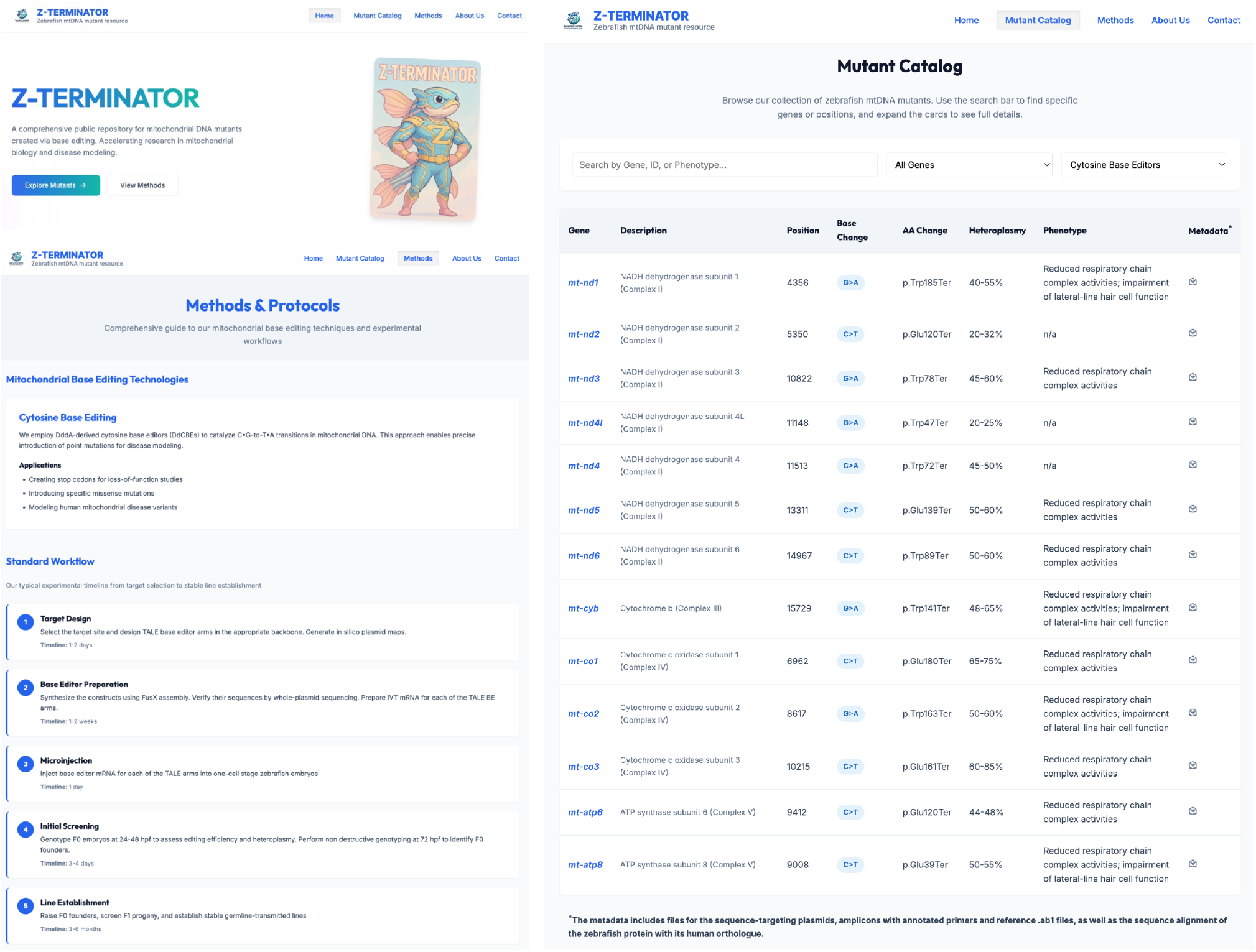
Z-Terminator - The Zebrafish mtDNA Mutant Resource. Screenshot of the resource homepage. The homepage summarizes the resource and links to the mutant catalog and methods section. The site provides open access to all reagents and sequencing metadata for the mutant collection described in this study.

## Discussion

Mutations in the mitochondrial genome, especially in protein-coding genes, produce distinct clinical syndromes. The canonical *MT-ND4* m.11778G>A mutation in Leber hereditary optic neuropathy (LHON)^75^ is typically homoplasmic or near-homoplasmic across tissues, yet it only affects retinal ganglion cells^76^, indicating cellular selectivity. This pattern underscores that MT-ND4, and likely other mtDNA-encoded Complex I subunits, have roles that extend beyond ATP production that are shaped by the developmental and metabolic context of individual cell types. These observations also highlight the need to examine the role of these genes beyond the bioenergetic lens, especially in a tissue and developmental context. MITOMAP catalogs hundreds of variants to which pathogenicity has been attributed^77^. However, vertebrate *in vivo* animal models harboring these variants have remained limited, largely due to the inability to introduce precise edits in the mitochondrial genome. In this study, we have established Z-Terminator, a vertebrate *in vivo* toolkit of loss-of-function alleles for the protein-coding genes of the mitochondrial genome. This library facilitates the interrogation of the role of individual OXPHOS subunits in mitochondrial function *in vivo*. Recent parallel efforts have extended the utility of DdCBEs to other vertebrate systems, to ablate all 13 protein-coding genes in mouse cell lines^49^, and conditional knockouts in rats using the Cre-lox system^50^. Extending this systematic strategy to zebrafish creates a complementary whole-organism model suited to pharmacological screening, real-time developmental imaging, and in *vivo* functional analysis.

To navigate the sequence constraints of the mitochondrial genome and achieve coverage at all 13 protein-coding loci, we adopted a dual-architecture design. Canonical DdCBEs require a 5’ T at the N-terminal TALE binding position, which precluded base editing at 6 of the 13 loci when these constrained designs were tested alone. To overcome this limitation, we incorporated unconstrained DdCBEs (also known as αDdCBEs) to our pipeline. These unconstrained base editors relax the canonical 5’ T requirement in the TALE N-terminal domain^35^, which enabled targeting at the otherwise inaccessible mtDNA loci. Notably, editing efficiencies were comparable between constrained and unconstrained architectures, indicating that the 5′-T preference reflects TALE-arm geometry rather than an intrinsic requirement of the deaminase.

To achieve DdCBE-mediated PTC installment, we leveraged two features of the vertebrate mitochondrial genetic code, both of which are central to the Z-Terminator designs. In vertebrate mitochondria, the UGA codon encodes tryptophan rather than a stop codon, and in-frame CAA encodes glutamine. A C-to-T transition at either position generates a UAA premature termination codon, converting a sense codon directly into a stop without relying on frameshift mutations or indel-based disruption. Expectedly, we observed locus-dependent variability in editing efficiency between the 13 tested loci. We attribute this to differences in local sequence context and the accessibility of individual target cytosines to DdCBEs. Despite low initial heteroplasmy at some loci, these edits can be enriched by base editor microinjection in founder animal progeny. Alternatively, these low-frequency founder edits may segregate to higher heteroplasmy levels in F1 progeny through the germline bottleneck.

We identified a cross-complex interdependency of bioenergetic phenotypes in our Z-Terminator library of mutant zebrafish lines. Notably, Complex II activities were unchanged across all genotypes, with only a slight decrease in *mt-co1* mutants, confirming the mtDNA specificity of these phenotypes. In contrast, Complex I activity was not only reduced in larvae harboring PTCs in Complex I subunits, but also in larvae with PTCs in subunits of Complex III, IV, and V. This observation, in which a defect in an mtDNA-encoded OXPHOS complex subunit secondarily affects the activity of other OXPHOS complexes, has been documented in patient-derived cell lines. In human 143B cybrid cells containing a m.3571dupC homoplasmic *MT-ND1* mutation, loss of functional MT-ND1 protein not only affects Complex I assembly but also reduces the steady-state levels of Complex IV^78^. This intercomplex regulation can be attributed to the diminished stability of the respiratory supercomplex^65,79^. In another study, Borlado *et al.* reported that fibroblasts derived from patient tissues harboring a homoplasmic m.15533A>G mutation in *MT-CYB* displayed combined deficiency in the activities of Complex I, III, and IV^80^. These findings support our observations on intercomplex regulation as a conserved feature of OXPHOS biogenesis in the vertebrate respirasome architecture. Defects in Complex III impair Complex I biogenesis by attenuating the incorporation of the NADH module^81^. The CO1-HIGD2A submodule normally stabilizes the I+III₂ supercomplex^82^ and Complex I and III form a stable core respirasome to which Complex IV can also bind. It has been shown that reduced COX1 levels can progressively destabilize Complex I^82–85^. For Complex V readouts, we observe decreased Complex I activity in our *mt-atp6* and *mt-atp8* mutants. The reduction maybe a readout of a bioenergetic feedback mechanism where loss of ATP synthase prevents the proton dissipation, which may lead to mitochondrial hyperpolarization activating the m-AAA protease AFG3L2, mediating the proteolytic breakdown of Complex I^86^. However, these hypotheses require further validation.

In *mt-nd1*, *mt-nd3* and *mt-nd6* mutants, we observed the divergent changes in Complex IV activity (no change, significantly reduced, and significantly increased, respectively), indicating altered supercomplex integrities and differential impacts on reactive oxygen species production. Additional studies will be required to elucidate the mechanisms underlying the complex-specific activity changes. Lactate was significantly elevated in the majority of the mtDNA mutant animals, confirming a shift toward anaerobic glycolysis consistent with impaired OXPHOS. Lactic acidosis is a clinically validated biomarker of mitochondrial disease^87^, and its recapitulation in zebrafish supports the utility of this model for studying OXPHOS deficiency *in vivo*.

Hearing loss is one of the most common clinical manifestations of mitochondrial disorders. Multiple mutations in mtDNA genes have been linked to auditory dysfunction^51,88^. In a cohort of patients with the m.3243A>G mutation, the majority presented with sensorineural hearing loss^87,89^. As a comorbidity, a subset of individuals have also been reported to exhibit hearing loss as a clinical feature overlapping with kidney-specific manifestations^90^. However, no study has directly demonstrated that engineered loss-of-function alleles in these genes impair hair cell mechanotransduction in a living vertebrate. The sensitivity of hair cells to OXPHOS dysfunction reflects their high energetic demand. Hair cells of the inner ear exhibit high mitochondrial volume with a distinct spatial organization, with small apically localized mitochondria and a more basally localized reticular network^69,91^. This sensitivity to OXPHOS disruption is supported by previous work describing that approximately 75% of the ATP of inner ear hair cells is generated by OXPHOS^92^.

Zebrafish have been proven to be an excellent model to study hair cell pathology and hearing loss^91^. Zebrafish lateral line hair cells are structurally and functionally analogous to mammalian inner ear hair cells. In recent years, zebrafish have gained significant attention, with multiple nuclear-encoded mitochondrial genes implicated in hearing loss^91,93–95^. We used the YO-PRO-1 vital dye uptake assay to measure hair cell activity in the lateral line neuromast^62^. In our Z-Terminator library, YO-PRO-1 uptake was significantly reduced in *mt-nd1*, m*t-cyb*, *mt-co2*, and *mt-atp8* mutant larvae compared to wild-type controls, affecting Complex I, Complex III, Complex IV, and Complex V, respectively. The YO-PRO-1 phenotype spanned alleles affecting four distinct complexes, suggesting that hair cell dysfunction reflects a complex-independent threshold response to OXPHOS capacity. This genotype-to-functional threshold was also observed in a subset of animals harboring PTC alleles particularly in *mt-cyb* mutants, where animals displaying greater than 70% heteroplasmy showed near-complete loss of YO-PRO-1 uptake. This pattern is clinically concordant with manifestations that become more severe as mtDNA mutant heteroplasmy increases. These findings support the use of zebrafish for the study of hearing pathophysiology and the design of therapeutic regimens for mitochondrial hearing disorders. However, further investigation is needed to delineate whether the inner ear hair cell dysfunction is linked to a decrease in hair cell number or defects in mechanotransduction channel activity.

Tissue-specific heteroplasmy profiles documented in the Z-Terminator library provided a direct *in vivo* readout of somatic mtDNA segregation and germline bottleneck dynamics. Systemic retention of DdCBE-engineered edits was observed across the six tissues profiled: brain, eye, heart, liver, caudal fin, and ovaries. For *mt-nd1*, *mt-nd5*, and *mt-atp8*, edits were retained in founder females into adulthood. Z-Terminator alleles are stably maintained across organ types throughout adult life. Brain, eye, and heart showed consistent heteroplasmy within individual fish across all three loci, suggesting that high-energy demand tissues may constrain mtDNA pool composition over time. Ovarian heteroplasmy was lower than somatic heteroplasmy in most founder females across all three loci. This can be explained by the mtDNA bottleneck^96^, a stochastic reduction in effective mtDNA copy number during postnatal folliculogenesis where a subset of the maternal mtDNA pool is transmitted to each oocyte^97^. As the size of this bottleneck varies among individuals, somatic mtDNA heteroplasmy was not a reliable predictor of F1 transmission frequency. This is consistent with other reports demonstrating that each oocyte samples a stochastic, non-representative fraction of the mtDNA pool^72,96,98–100^. Two mtDNA founder females with comparable somatic heteroplasmy in the *mt-nd1* locus produced F1 clutches with divergent distributions. One female transmitted edits uniformly across sampled embryos, whereas the other produced a clutch in which most embryos carried less than 10% mutant mtDNA. This divergence from common somatic mtDNA heteroplasmy can be explained by inter-individual variation in bottleneck size as described above. This germline data illustrates the bottleneck constraints for the establishment of stable lines for these alleles. The *mt-atp8* locus showed the most pronounced amplification, where a founder female with 37% somatic heteroplasmy produced F1 embryos displaying up to 80% mutant mtDNA. The locus-dependent difference in transmission variance likely reflects purifying selection during the germline bottleneck^99,101^. Our data are consistent with stochastic variance during the germline bottleneck, though a larger sample size is required to investigate whether this variance is locus dependent.

In summary, we present Z-Terminator as a comprehensive mitochondrial base editing toolkit and *in vivo* atlas that enables systematic loss-of-function interrogation of all 13 mtDNA-encoded OXPHOS subunits in a vertebrate model organism. The Z-Terminator atlas establishes a genotype-resolved framework and offers a unique opportunity to serve as a functional complement to variant databases such as MITOMAP. All Z-Terminator sequence targeting reagents are publicly available, providing the mitochondrial disease community with a genotype-defined, whole-organism resource for dissecting tissue-specific consequences of mtDNA dysfunction, pharmacological screening, and therapeutic development.

## Acknowledgements

The authors thank Dr. Steven A. Farber and Monica Hensley (Johns Hopkins University, USA), Dr. Matthias Hammerschmidt and Dr. Joy Armistead (University of Cologne, Germany), Dr. Lisa A. Schimmenti and Joseph A. Dugdale (Mayo Clinic), and Dr. Marni J. Falk and Dr. Christoph Seiler (The Children’s Hospital of Philadelphia, USA) for providing valuable input during the preparation of this article. The authors also thank the former and current Ekker lab members Bibekananda Kar, Kavini Nanayakkara, Daniel M. Chambers, and Anjana Reddy, as well as the zebrafish facility staff at Mayo Clinic and The University of Texas at Austin.

## Authors’ Contributions

The idea was conceived by A.S. and S.C.E. The article was written by A.S., K.S.M, G.K.V and S.C.E. Mitochondrial DNA mutant lines using constrained editors were generated by A.S., K.S., and L.R.T. Mitochondrial DNA mutant lines using unconstrained editors were generated by M.R.A. J.K, and A.S. Experiments were executed by A.S., K.S.M., M.R.A., J.M., J.K., R.S., K.S., L.R.T., A.Z., C.P., and M.M. with experimental guidance from G.K.V., E.N.O., K.J.C., and S.C.E. Data analysis was carried out by A.S., K.S.M., M.R.A, J.K., G.K.V., and E.N.O. The unconstrained base editor backbone for generation of zebrafish constructs was provided by S.R.C. The article was reviewed and edited by A.S., K.S.M., M.R.A., J.K., L.R.T., S.R.C., M.M., G.K.V., E.N.O., and S.C.E.

## Data Availability

Plasmids are available upon request. The FusX assembly kit is available through Addgene (Kit no. 1000000063). Metadata for all sequence targeting reagents will be released upon publication.

## Author Disclosure Statement

The Mayo Clinic has an issued patent (US20180002707A1) on the FUSX TALE assembly system used here. S.C.E is a shareholder in LifEngine Technologies that has a license to the FUSX TALE assembly system.

## Funding Information

This work was funded by grants from the NIH GM063904, NIH R24OD035577, Dell Medical School at The University of Texas at Austin, Mayo Foundation for Medical Education and Research, and The Children’s Hospital of Philadelphia (CHOP) Mitochondrial Medicine Frontier program. G.K.V and C.P are supported in part by funding from the NIH/ORIP (R24OD034438).

## Supplementary Material

### Supplementary Tables

**Supplementary table 1:**
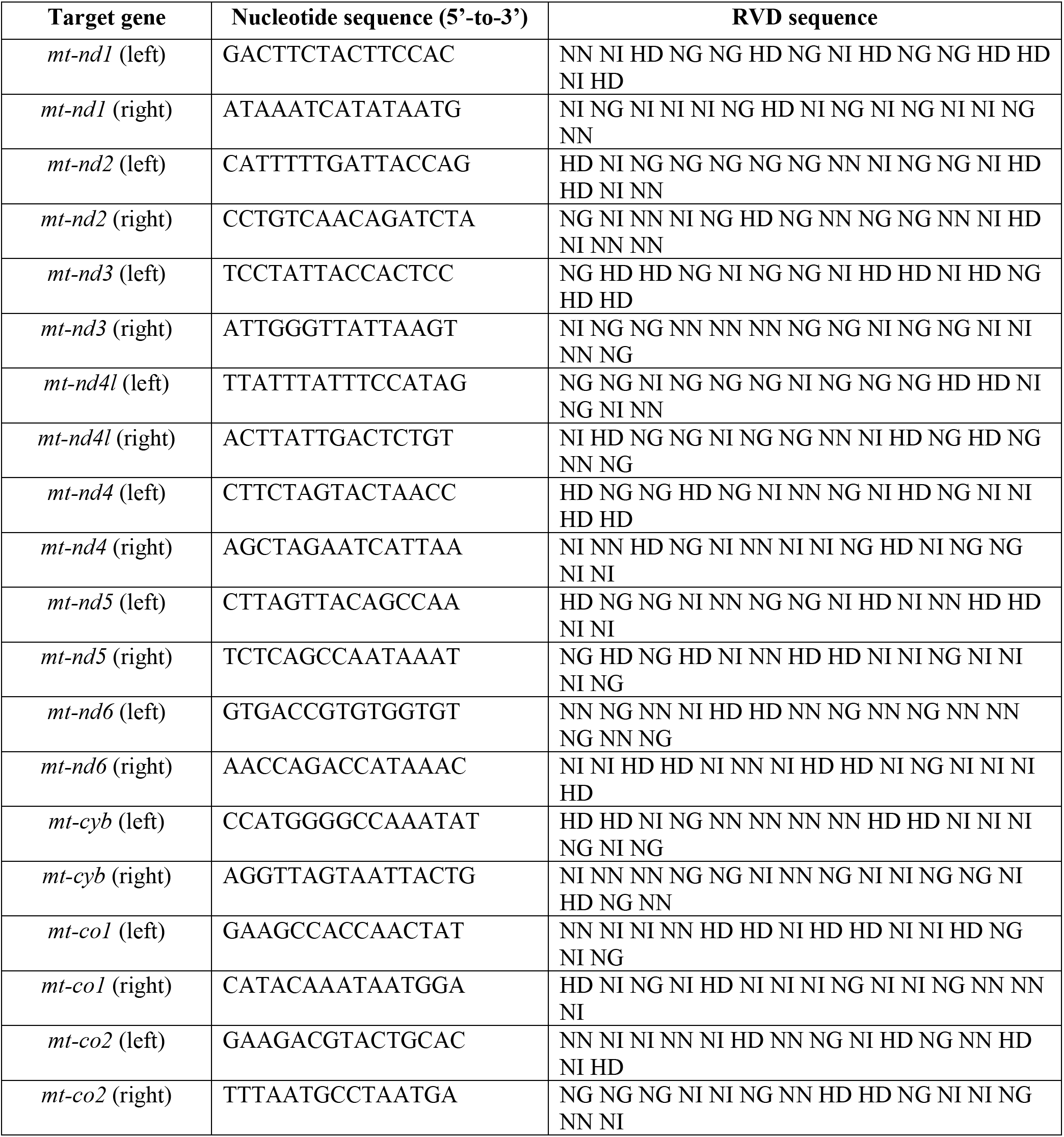
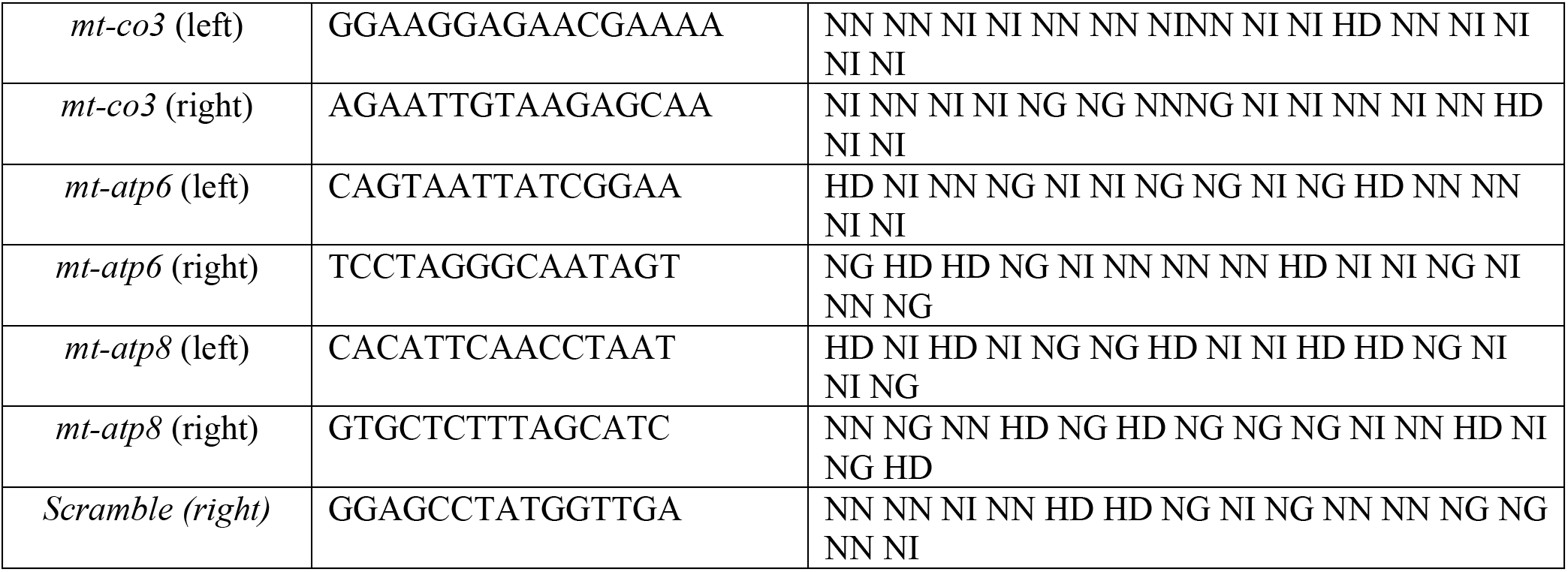
RVD sequences for mtDNA loci targeted in Z-Terminator.

**Supplementary table 2:**
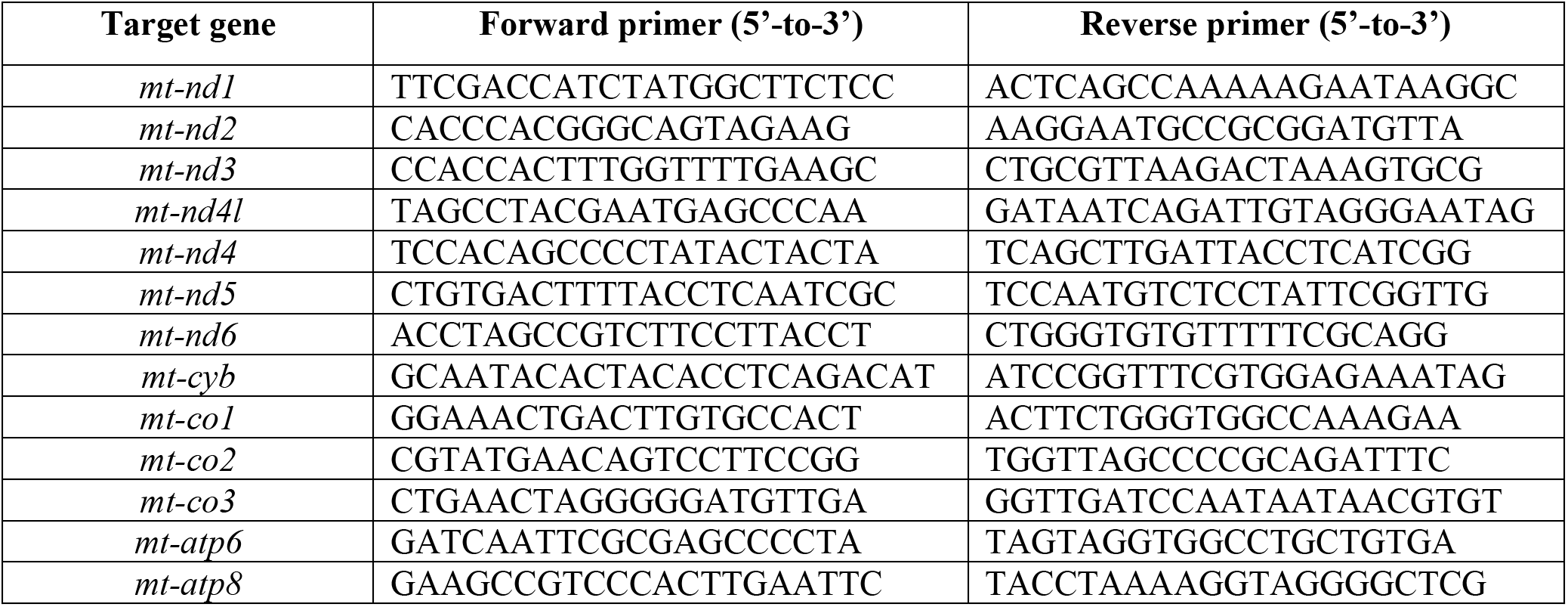
List of oligonucleotides used in this study.

### Supplementary Figures

**Supplementary Figure 1:**
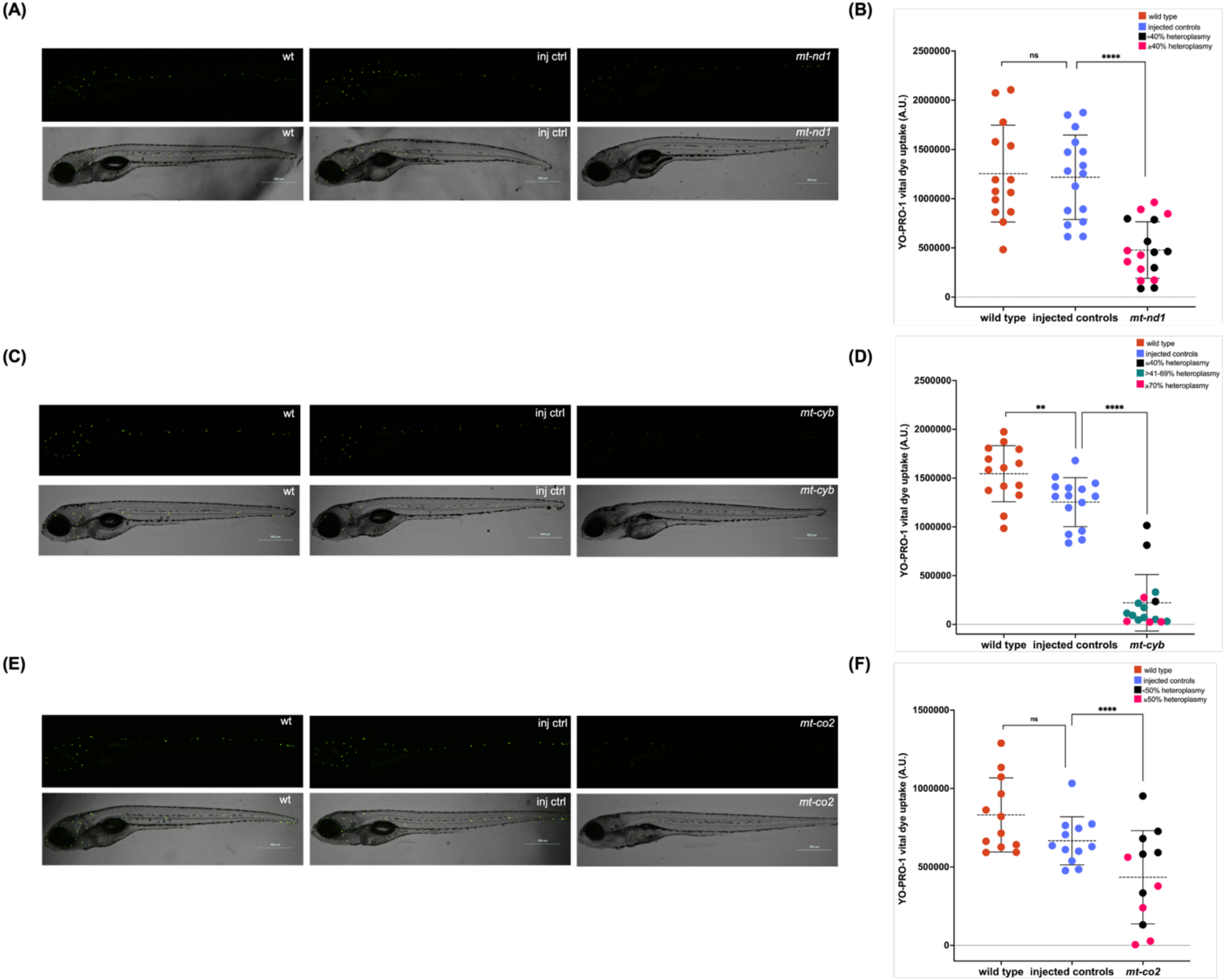
Reduction of YO-PRO-1 dye uptake in mtDNA mutant larvae is attributable to the engineered PTC allele and not to the microinjection procedure. **(A, C, E)** Representative YO-PRO-1 fluorescence and corresponding brightfield images of wild-type, injected-control, and mutant larvae at 5 dpf for *mt-nd1* **(A)** *mt-cyb* **(C)**, and *mt-co2* **(E)**. Mutant larvae show a reduction in neuromast YO-PRO-1 labeling relative to both wild-type and injected-control groups. Magnification-4X; Scale-bar 500 μm. **(B, D, F)** Corrected total fluorescence (CTF) of YO-PRO-1 vital dye uptake at the MI1 neuromast in wild-type, injected-control, and mutant larvae for *mt-nd1* **(A)**, *mt-cyb* **(C)**, *mt-co2* **(E)**. Each data point represents an individual larva. p-values for pairwise comparisons were determined by Student’s t-test (**p < 0.01; ***p < 0.001; ****p < 0.0001). Horizontal lines indicate the mean and error bars represent the standard deviation. dpf, days post-fertilization. wt-wild type; inj ctrl-injected control.

**Supplementary Figure 2:**
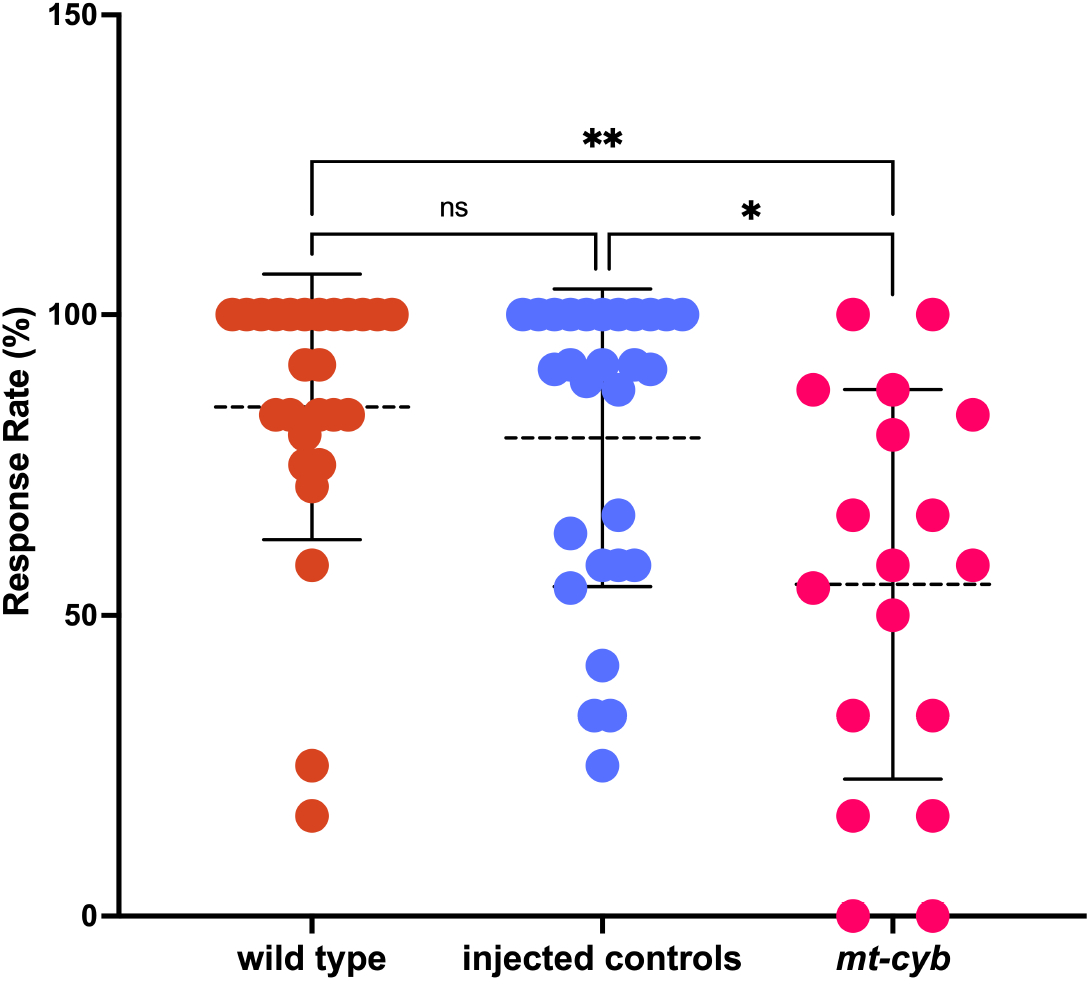
*mt-cyb* mutant larvae show a reduced acoustic-evoked behavioral response. Larvae were assayed for AEBR at 6 dpf. Response rate was comparable between tab5 wild-type and injected control larvae. *mt-cyb* mutant larvae showed a significantly reduced response rate compared with both tab5 (p < 0.01) and injected control groups (p < 0.05). Each data point represents an individual larva. Statistical significance was determined by Kruskal-Wallis test. Horizontal lines indicate the mean and error bars represent the standard deviation. dpf, days post-fertilization.

